# Epithelial eversion, a collective rearrangement from apical-in to apical-out polarity, is initiated by α6β4 integrins and sustained by increased cell proliferation and anchorage-independence

**DOI:** 10.1101/2025.05.12.653281

**Authors:** Niharika Patel, Jenna Pirrello, Shivam Priya, Aki Manninen, Daniel Conway

## Abstract

Epithelial cells primarily segregate transmembrane proteins to apical or basal surfaces, establishing apical-basal polarity. For 3D tissues, apical proteins face inwards and basal proteins are on the outer surfaces (in contact with ECM). Apical-basal polarity can become ‘inverted’ such that the apical proteins are on the outer surfaces. Originally this polarity inversion was assumed to always occur due to changes in protein trafficking. However, recent work by our group showed that increased RhoA activation causes epithelial spheroids to invert apical-basal polarity via a collective rearrangement of cells, a process we and others have termed eversion. We hypothesized that specific integrin-ECM interactions could facilitate the collective cellular migration required for eversion. Using multiple integrin knockout lines, we observed that epithelial cells lacking either α6 or β4 integrins do not develop apical-out polarity in response to RhoA activation. Similarly, α6 blocking antibody or culturing spheroids in collagen (lacking laminin) also inhibited the formation apical-out polarity. These data indicate that α6ß4-laminin interactions are required for eversion. Next, we examined the role of cell proliferation and anchorage independence in maintaining apical-out polarity of everted spheroids. Inhibition of cell proliferation (with DNA synthesis inhibitor aphidicolin) or inhibition of anchorage-independence (with FAK inhibitor 14) were sufficient to restore apical-in polarity to RhoA treated spheroids, indicating that both proliferation and anchorage-independence maintain spheroids in an apical-out state. We also observed that apical-out spheroids can ‘revert’ to apical-in polarity through apoptotic cavitation of cells located in the center of the spheroids. Lastly, through RNA sequencing we demonstrate that apical-out spheroids have unique gene expression profiles. This study provides new mechanistic insights into the biochemical and biophysical mechanisms that drive eversion and maintain apical-out polarity. This work supports the concept that changing from apical-in to apical-out polarity be an important marker for phenotypic switch in epithelial cells.

## Introduction

Establishment and maintenance of apical-basal polarity is a critical function for epithelia that develops and maintains epithelial tissue organization.^1^ This involves the segregation of transmembrane proteins into distinct vesicles at the trans-Golgi network for trafficking to either the apical or basal sides of the cell.^2^ In ductal, glandular, or tubular epithelial tissue structures apical-basal polarity is established such that the apical membrane surface faces inwards (apical-in) towards a liquid-filled lumen structure. The formation and maintenance of the lumen structure is driven by establishment of apical-basal polarity. Apical actin polymerization has been shown to drive the initial formation of a lumen.^3^ Following the establishment of barrier function through tight junction formation, apical ion secretion establishes an osmotic pressure that mechanically stabilizes the lumen.^3–5^ In addition to the role of apical actin polymerization and osmotic pressure for driving lumen formation, under certain conditions apoptotic cavitation has also been shown to be important factor for the formation of lumens.^6^

Loss of apical-basal polarity can drive tissue disorganization and is a hallmark of carcinoma (cancers originating from epithelia).^7^ In addition to a complete disruption of apical-basal polarity, it is also possible for apical-basal polarity to become ‘inverted’ where apical and basal transmembrane proteins remain segregated but instead are organized such that the apical surface faces outwards (apical-out). Apical-out polarity is a defining characteristic of specific epithelial cancers, including micropapillary carcinomas, and is associated with increased invasion.^8^ Mostov and colleges have shown that apical-out polarity can be established by various stimuli that alter apical-basal protein-trafficking.^9,10^

Recently our group and others have shown that epithelial spheroids can transition from normal, apical-in polarity to apical-out polarity through a collective rearrangement of cells that flip the spheroid inside-out, a process that we and others have termed eversion.^11–13^ Our group developed a biophysical model to explain eversion, in which differences in the apical and basal surface energies provides a driving force for eversion.^11^ While others have observed eversion when spheroids are removed from ECM,^12,13^ a notable aspect of our prior work was that we were able to observe eversion of spheroids within ECM,^11^ which suggests the possibility that eversion could occur in physiological or pathological conditions. Despite our biophysical model for how mechanical forces promote eversion, there is still an incomplete understating of the biochemical mechanisms that promote eversion and sustain apical-out polarity.

Given the collective movement of cells during eversion, we speculated that specific integrin-ECM interactions may be necessary for eversion to occur. By screening the response of integrin knockout MDCK cell lines to an activator of Rho, we determined that cells lacking α6 or β4 integrin had significantly reduced incidence of eversion, even though these cells exhibited lumen collapse in response to Rho activation. α6 blocking antibodies also inhibited RhoA-driven eversion in both MDCK and Caco-2 cells. Additionally, MDCK spheroids cultured in collagen failed to assemble a basal lamin-ECM network and did not evert in response to RhoA activation. These data indicate that α6β4-laminin interactions are likely necessary for enabling the collective movement of cells required during eversion to invert apical-basal polarity. We also show that everted, apical-out spheroids can return to apical-in polarity when RhoA activator is removed by undergoing apoptotic cavitation. Both increased proliferation and anchorage independence are required for maintaining apical-out polarity, as inhibitors of DNA synthesis and focal adhesion kinase activity prevent eversion and restore normal apical-basal polarity. Lastly, through RNA sequencing we demonstrate that apical-out spheroids have unique gene expression profiles. In summary, our study provides new mechanistic insights into the biochemical and biophysical mechanisms that drive eversion and maintain apical-out polarity.

## Methods

### Cell lines and culture

MDCK II cells, originally a gift from Rob Tombes (Virginia Commonwealth University), and Caco-2 (HTB-37, ATCC) were used in these studies. Additionally, to examine the role of specific integrins in the eversion process, we utilized a set of existing α3, αV, α6, β1, β4, β1 and β4 integrin knockout cells that were previously established from the MDCK-II Heidelberg strain.^14–16^ In experiments using integrin knockout cells they were exclusively compared to the parental MDCK-II Heidelberg strain. All cell lines were cultured in DMEM medium with 4.5 g/L glucose (ThermoFisher), supplemented with 10% v/v fetal bovine serum (FBS, ThermoFisher) and 1% v/v penicillin-streptomycin mix (Pen-Strep, ThermoFisher).

### EdU incorporation assay

*For proliferation*, Click-iT™ Plus EdU Cell Proliferation Kit for Imaging, Alexa Fluor 488 (ThermoFisher catalog C10637) was used. Edu was added to the cells 24 hours prior to fixation. The next day the cells were fixed and stained per manufacturer instructions.

### Inhibition experiments

Aphidicolin was used at a final concentration of 2 μg/mL (Sigma Aldrich catalog A4487) to inhibit DNA synthesis. Focal adhesion kinase (FAK) inhibitor Y15 (also known as FAK inhibitior 14) was used at a final concentration of 2.5 μM (Medchem express, catalog HY-12444).

### Formation and culture of MDCK spheroids in Matrigel

For formation and culture of 3D spheroids in Matrigel, cells were trypsinized from tissue culture plates and 5000 cells were suspended in 400 μL of growth medium supplemented with 2% v/v Matrigel. The cell suspension was seeded onto an eight-well Nunc Lab-Tek II chambered coverglass or coverslip, pre-coated with 40 μL of phenol-free growth-factor-reduced (GFR) Matrigel (Corning) as previously described.^17^ The pre-coated Matrigel was allowed to solidify by incubation for at least 30 min at 37°C before seeding the cells. The growth medium was changed every 3 to 5 days and the cells were allowed to form acini for 7–12 days. To induce eversion spheroids were treated with Rho activator II (Cytoskeleton) at a final concentrations of 2-4 μg/mL for 48 h. In some batches of Matrigel lower concentrations (1-2 μg/ml) of Rho activator II did not induce eversion. Because of inconsistencies with eversion with different lot numbers of Matrigel, the Rho II activator concentration was chosen such that eversion consistently occurred in wild-type, untreated cells without any evidence of loss of cell-cell cohesion (loss of cell-cell adhesion was previously observed at doses of 5 μg/mL or 10 μg/mL Rho activator II^11^).

### Formation and culture of MDCK spheroids in collagen

For formation and culture of 3D spheroids in collagen, cells were trypsinized from tissue culture plates and suspended in a collagen type 1 solution (PureCol catalog#5005, Advanced BioMatrix) and seeded onto eight-well Nunc Lab-Tek II chambered coverglass as previously described.^18^ Briefly, collagen solution was prepared by mixing 50 μL GlutaMax (200 mM), 625 μL NaHCO□ (2.35 mg/mL), 625 μL MEM 10X, 125 μL HEPES (1M, pH 7.6), and 4.13 mL Collagen 1 (2 mg/mL) to a final volume of 6.255 mL at neutral pH. Eight-well chambers were coated with the collagen solution with 40ul and incubated for polymerization in 37°C oven for 30 minutes. Cells were diluted into the collagen solution at 10,000 cells/mL, ensuring the cell volume does not exceed 10% of the total solution. 40 μL of the cell-collagen mixture was applied to pre-coated plates. The growth medium was changed every 3 to 5 days and the cells were allowed to form acini for 7–12 days. To determine if spheroids cultured in collagen underwent eversion spheroids were treated with Rho activator II (Cytoskeleton) at a final concentration of 3.5 μg/mL for 48 h.

### Formation and culture of Caco-2 spheroids

Caco-2 spheroids were formed similarly to MDCK spheroids, except that a mixture Matrigel and Collagen was used (collagen I, final concentration of 1 mg/ml; Matrigel, final concentration of 40% or more.^19^ On 6^th^ day they were treated with cholera toxin (0.1ug/ml, Sigma) to enhance lumen formation. ^19^

### α6 blocking experiments

Wild-type (WT) cells were cultured in Matrigel within eight-well chambers. After 7–10 days of culture, the resulting acini were exposed to different experimental conditions. The control wells remained untreated, while others were treated with Rho Activator II, or Rho Activator II combined with the α6 functional blocking antibody (clone GoH3, BioLegend) at a concentration of 10 μg/mL. Following 48 hours of treatment, the cells were fixed using 2% paraformaldehyde (PFA).

### Immunofluorescence

For immunofluorescent staining experiments cells were fixed in 2% paraformaldehyde, followed by 0.1 % Triton X permeabilization. Primary antibodies used were podocalyxin/gp135 (clone 3F2:D8, Sigma), α6 integrin (clone GoH3, BioLegend), β4 integrin (clone M126, Abcam), laminin (catalog:L9393, Sigma Aldrich), and phospho-Ezrin/Radixin/Moesin (catalog: 3726S, Cell Signaling). Alexa Fluor secondary antibodies (ThermoFisher Scientific) were used. Actin was labeled using fluorescently tagged phalloidin (Cytoskeleton). Nuclei were labeled with Hoechst from Thermofisher (catalog: H3570). Cells were mounded with ProLong Gold Antifade Mountant (ThermoFisher Scientific catalog: P36930) and imaged on a Stellaris 8 confocal microscope (Leica).

### Tunnel apoptosis assay

Wild-type (WT) MDCK cells were seeded into Matrigel and cultured in eight-well Nunc Lab-Tek II chambered cover glasses for 7-12 days. Acini formation was observed within 7–10 days. Subsequently, the acini were treated with Rho Activator II for 48 hours and fixed at various time points following the removal of Rho Activator II, specifically at 0, 24, 48, and 96 hours. Cells were fixed using paraformaldehyde (PFA) and apoptosis was assessed using the Click-iT TUNEL Assay Kit (C10619, Thermo Fisher Scientific) per manufacturer instructions. To evaluate the apoptotic effects of the α6 blocking antibody, WT cells were subjected to the following experimental conditions: a control group without Rho Activator II, a group treated with Rho Activator II, and a group treated with Rho Activator II in combination with the α6 blocking antibody. Cells were fixed using paraformaldehyde (PFA) and apoptosis was assessed using the Click-iT TUNEL Assay Kit (C10619, Thermo Fisher Scientific).

### Bulk RNAseq experiments

MDCK cells were seeded into Matrigel in 12 well plates at 11,000 cells per well. WT and α6 KO cells were allowed to form spheroids for 7-12 days, followed by 48 hours of Rho Activator II treatment. Cells were harvested from matrigel by dissolving matrigel using Corning cell recovery solution (catalog: 354253). Trizol (ThermoFisher Scientific) was used to extract RNA as per manufacturer protocol. The extracted RNA was then purified using the NEB Clean-Up Kit (catalog: T2030). Purified RNA was quantified using a NanoDrop spectrometer.

The RNA was sent to Novagene (Sacramento, CA) for RNA seq analysis. Messenger RNA was purified from total RNA using poly-T oligo-attached magnetic beads. After fragmentation, the first strand cDNA was synthesized using random hexamer primers, followed by the second strand cDNA synthesis using either dUTP for directional library or dTTP for non-directional library. The library was checked with Qubit and real-time PCR for quantification and bioanalyzer for size distribution detection. Quantified libraries will be pooled and sequenced on Illumina platforms, according to effective library concentration and data amount. Raw FASTQ reads are processed with fastp to remove adapters, poly-N sequences, and low-quality reads. Clean reads are aligned to the reference genome using Hisat2, and novel transcripts are predicted with StringTie. Gene expression levels are quantified using FeatureCounts, and differential expression is analyzed with DESeq2 or edgeR. Functional enrichment analyses, including GO, KEGG, and GSEA, were performed using the cluster Profiler R package. RNAseq data was deposited at NCBI under accession PRJNA1260979.

### Agar assay

To assess anchorage-independence a standard agar assay was used.^20^ Wild-type (WT) cells were seeded in T25 flasks two days before the assay. One group remained untreated as the control, while the experimental group was pre-treated with Rho Activator II at a concentration of 2 μg/μL. The day prior to the experiment, six wells were coated with a 1% F127 solution in PBS for one hour to prevent cell attachment to the surface. Agarose (Fisher Scientific catalog: BP-165-25) solutions at concentrations of 4% and 1.75% were prepared in ultrapure water and media containing 12% FBS was also prepared. To create the bottom layer, the 4% agarose solution was mixed with media in a 1:8 ratio, and 2 mL of this mixture was used to coat each well. The plates were allowed to set for one hour in the biosafety hood. Cells were detached from T25 flasks by trypsin, counted, and approximately 15,000 cells were added per well. For the top layer, 8 mL of cell-suspended media was combined with 2 mL of 1.75% agarose solution. A total of 2 mL of this mixture was applied on top of the bottom layer in each well and allowed to solidify for an additional three hours. The following day, 1 mL of media was added to each well every week, with the experimental group receiving media supplemented with Rho Activator II at a final concentration of 2 μg/μL (Rho activator was maintained during the 4-5 weeks of the agar assay). After 4-5 weeks colonies were visible, and images were taken with optical microscope.

### Statistics

For each spheroid eversion experiment, 10 spheroids per condition were scored as normal, lumen collapse, or eversion through the assessment of apical markers. Repeat experiments (on different days) were performed. Statistical differences were assessed using *p<0.05 as per ANOVA with Tukey’s multiple comparison test. For Agar Assay, Paired T test was performed.

## Results

### Involvement of α6β4 integrins and laminin ECM in eversion

Our previous studies of RhoA-driven changes in apical-basal polarity showed that eversion is a collective process that requires cell-cell adhesions.^11^ To test the involvement of integrins in the eversion process we assessed the ability of a set of existing α3, αV, α6, β1, β4, and β1+β4 integrin knockout MDCK cell lines^14–16^ to undergo RhoA-driven eversion. Apical polarity of MDCK spheroids was assessed using both actin and podocalyxin (gp135) immunostaining (**Figure 1A, Supplemental Figure 1**). α6 and β4 exhibited lumen collapse following Rho Activator II treatment (**Figure 1A, 1B**), indicating that lumen collapse is not dependent on these integrins. However, α6 and β4 knockout cell lines exhibited significantly reduced incidences of apical-out polarity following 48 hours of Rho Activator II treatment (**Figure 1A, 1B**), indicating that these integrins are important in the establishment of apical-out polarity. These data suggest that lumen collapse precedes eversion and that lumen collapse is not sufficient to promote eversion. β1 knockout and β1+β4 double knockout spheroids exhibited apical-out polarity before Rho Activator II treatment (**Supplemental Figure 1A**), which is consistent with prior reports showing apical-out polarity with β1 integrin blocking antibodies.^23^ Because β1 and β1+β4 knockout cells already had apical-out polarity we did not explore the effects of Rho activation. α3 knockout cells, which exhibited a previously described multiple lumen phenotype^21^ had some instances of apical-in and apical-out polarity, and did not consistently evert when treated with Rho Activator II (**Supplemental Figure 1B**). αV cysts also had a multiple lumen phenotype, which may relate to the altered mechanotransduction and focal adhesion previously seen with loss of αV expression.^22^ αV knockout cells still transitioned to apical-out polarity (**Supplemental Figure 1B**), indicating that αV is not required for eversion. Given the complex phenotypes of α3, αV, β1, and β1+β4 knockout lines, including disruptions in morphology prior to RhoA activation, these lines were not used for further studies.

**Figure 1.**
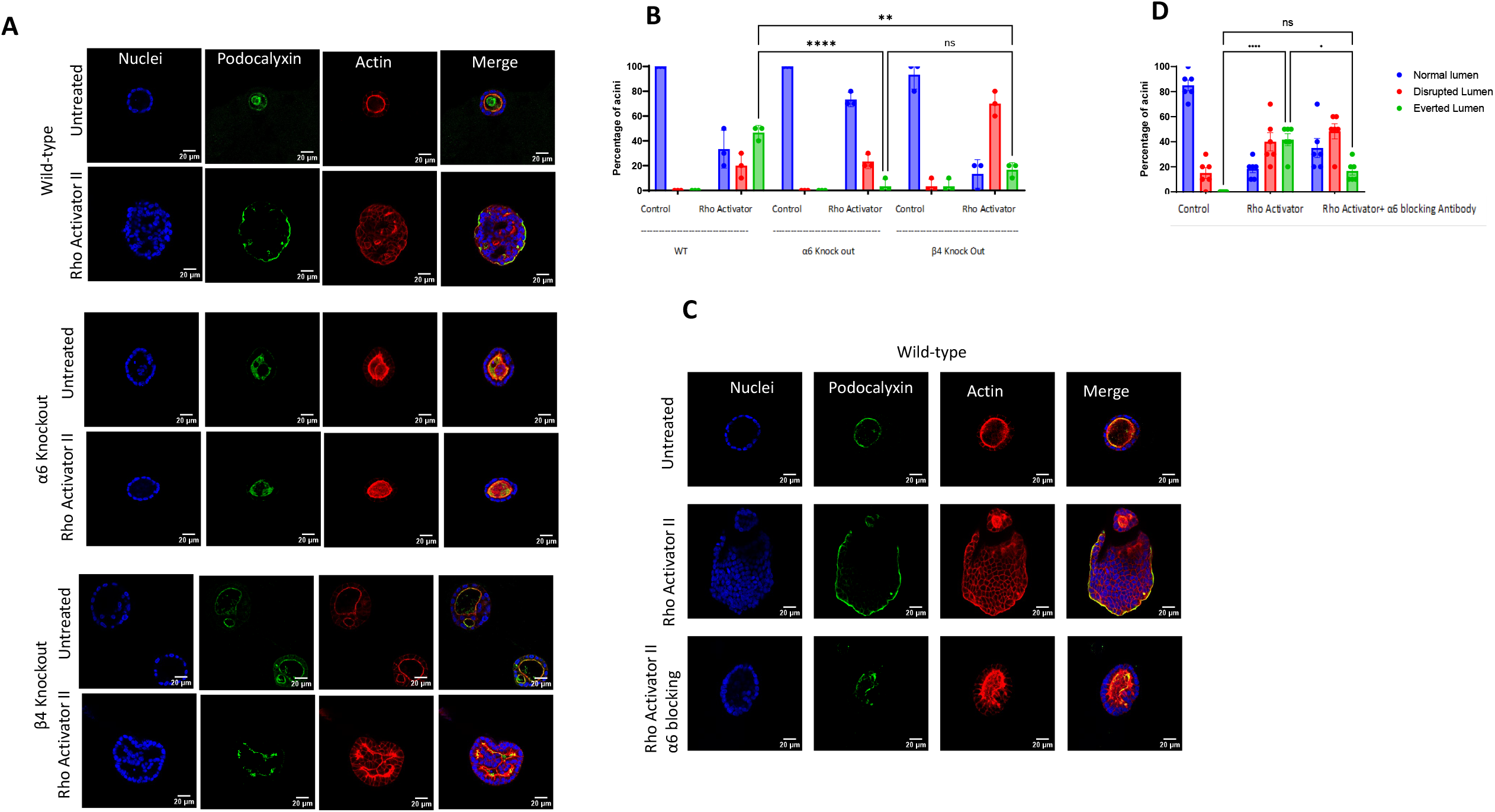
Loss of α6 or β4 integrins significantly inhibits Rho-A mediated eversion. **A)** MDCK cysts were formed by seeding wild-type, α6 integrin knockout, or β4 integrin knockout MDCK cell lines in Matrigel. MDCK cysts were then treated with RhoA activator II for 48 hours. Immunostaining was performed for podocalyxin and actin (phalloidin) to assess polarity **B)** Cysts were classified as normal (hollow cell-free lumen and normal apical-in polarity), disrupted lumen (loss of hollow lumen but normal apical-in polarity), or everted (lumen collapse and apical markers on the outer membrane of the cyst). The incidence of RhoA activator-induced eversion was significantly less in both α6 and β4 knockout cysts. **C)** MDCK cysts were left untreated, or treated with α6 blocking antibody, RhoA activator II, or α6 blocking antibody with RhoA activator II and then fixed and immunostained. **D)** Quantification of normal, disrupted lumen, and eversion for blocking antibody experiments.

To further test the importance of α6 integrins for eversion, we used the GoH3 functional α6 blocking antibody.^24^ Treatment of spheroids with α6 blocking antibody had minimal effects on apical-basal polarity and did not disrupt single lumen morphology (**Supplemental Figure 2**). However, α6 blocking antibody significantly inhibited the development of apical-out polarity for Rho Activator II treated wild-type MDCK cells with (**Figure 1C, 1D**). We repeated these experiments using Caco-2 cells, a human epithelial cell line which also assembles into spheroids with apical-in polarity. First, we confirmed that Caco-2 cells, when treated with Rho Activator II, developed apical-out polarity (**Supplemental Figure 3A, 3B**). Next, we showed that treatment with α6 blocking antibody also inhibited apical-out polarity by Rho Activator II in Caco-2 cells (**Supplemental Figure 3A, 3B**).

Because α6 and β4 integrins are important in the formation of laminin-binding hemidesmosomes, we sought to understand if there would be differences for cells grown in Matrigel versus collagen gels. Matrigel contains laminin whereas the collagen used does not any additional ECM proteins. We compared the formation of laminin ECM for MDCK cells grown in Matrigel vs collagen I ECM. MDCK spheroids formed a hollow lumen with correct apical-basal polarity in both Matrigel and collagen. When laminin ECM was assessed using a laminin antibody, spheroids cultured in Matrigel had strong immunostaining for laminin at the basal (outer) surface (**Figure 2A**). In contrast, spheroids cultured in collagen had a mixed phenotype in which the laminin network was either observed on the basal (outer) side of the spheroid or instead as assembled laminin at the apical surface (**Figure 2A**). Regardless of the location of laminin, we observed much weaker immunostaining of laminin for cells cultured in collagen as compared to Matrigel (**Figure 2A**). Prior reports have shown assembly of laminin on the basal surface for MDCK in collagen gels,^9,25^ although these prior studies did not directly compare laminin assembly for spheroids in collagen versus Matrigel. The reason for the inconsistent assembly of laminin at the basal (outer) surface of cysts in our collagen gels is not clear but may possibly involve different sources of collagen. Nevertheless, the reduced basal laminin assembly in collagen ECM provided us with the opportunity to explore how this would affect eversion. We treated spheroids in Matrigel or collagen gels with Rho Activator II (**Figure 2B**). Although spheroids in collagen exhibited a disrupted lumen structure, we were not able to find any instances of apical-out polarity for spheroids in collagen treated with Rho Activator II (**Figure 2B, 2C**). We also tried to determine if collagen-only culture of Caco2 cells would suppress Rho-driven eversion; however, Caco2 cells did not properly develop into spheroids when cultured in pure collagen (data not shown). Overall, our results in **Figures 1** and **Figure 2** show that α6β4-laminin interactions are necessary for establishment of apical-out polarity.

**Figure 2.**
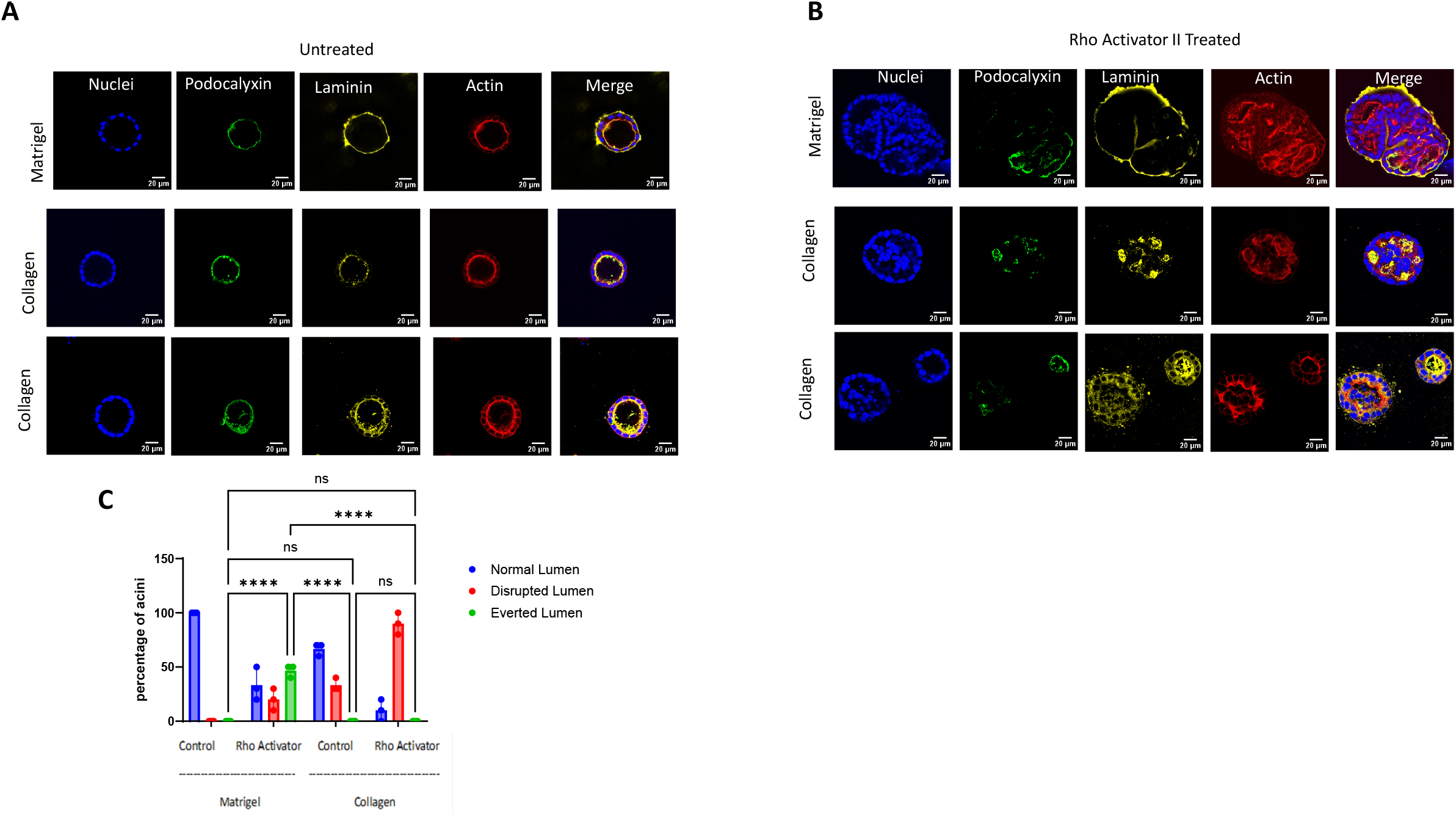
Cells do not evert in collagen. **A)** MDCK cells were seeded into Matrigel or Collagen allowed to form cysts. Cysts were then fixed and immunostained for podocalyxin, laminin, and actin (phalloidin). Less laminin basal laminin staining was observed for spheroids in collagen. **B)** MDCK cysts were treated with 48 hours of RhoA activator II and then fixed and immunostained for podocalyxin, laminin, and actin (phalloidin). **C)** Quantification of normal, disrupted lumen, and eversion for each condition.

### Apical-out spheroids can return to apical-in polarity through apoptotic cavitation

We sought to understand if eversion (changing from apical-in to apical-out polarity) is a reversible process. Spheroids were treated for 48 hours with Rho Activator II to induce eversion and then media was replaced with fresh media (without Rho Activator II). After 24 hours there was a statistical decrease in the number of spheroid with apical-out polarity and significant decrease in lumen disruption observed at 48 hours (**Figure 3A, 3B**). These data indicate that reversion is a reversible process. Following removal of Rho Activator II, we also observed altered nuclear morphologies for cells in the center of the spheroids (**Figure 3A**) that were suggestive of apoptosis. To assess apoptosis we used the TUNEL assay, which detects fragmented DNA in cells. Following removal of Rho Activator II, we observed an increase in apoptotic cells (**Figure 3C**). Notably these apoptotic cells were only observed at the center of the spheroid, and no apoptosis was observed for cells in the outermost layer in contact with ECM (**Figure 3C**). We also examined if treatment with α6 blocking antibody was sufficient to restore a hollow lumen. Spheroids were first treated with RhoA Activator II, followed by treatment with α6 blocking antibody (in the presence of RhoA Activator II). Spheroids treated with α6 blocking antibody also exhibited increased apoptosis at the center of the spheroid (**Figure 3D**). These data indicate that reversion is a reversible process, that hollow lumens are re-established by apoptotic cavitation, and blocking α6 integrins is sufficient to stimulate apoptotic cavitation.

**Figure 3.**
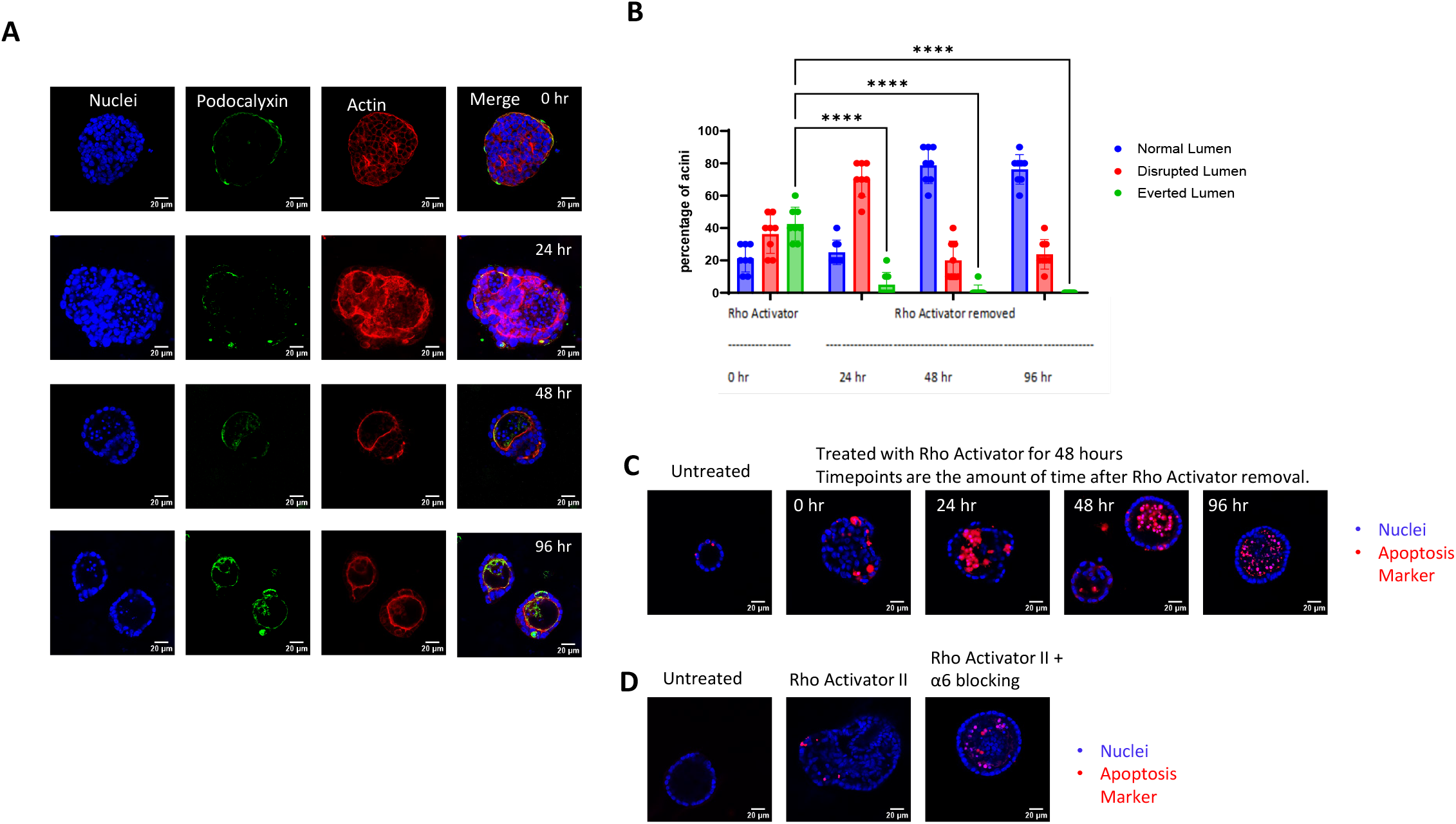
Reversion of apical-out polarity to apical-in polarity occurs through apoptotic cavitation. **A)** MDCK cysts were grown in Matrigel for 7 days, and then treated with RhoA activator II for an addition 48 hours. RhoA activator II was removed from culture by changing media. Cysts were fixed and immunostained at the indicated time points where time indicates the amount of time after RhoA activator removal. **B)** Quantification of normal, disrupted lumen, and eversion for each timepoint. **C)** Apoptosis was assessed using TUNEL assay at each timepoint. **D)** MDCK cysts were left untreated, treated with RhoA activator II, or RhoA activator II and α6 blocking antibody and apoptosis was assessed with the TUNEL assay.

### Apical-out polarity is maintained by anchorage-independence

Since we had observed that transitions from apical-out to apical-in polarity occur through apoptosis, we hypothesized that apical-out spheroids may exist by cells developing anchorage-independence (resistance to anoikis). To directly measure anchorage-independent cell growth we used a soft agar colony forming assay. Rho Activator II treated cells exhibited increased number and size of colonies in soft agar (**Figure 4A, 4B**). Next, we treated spheroids with an inhibitor of focal adhesion kinase (FAK) as inhibition of FAK has been shown to restore anchorage-dependence.^26^ Pre-treatment of spheroids with FAK inhibitor prevented infilling of the lumen with cells (**Figure 4C, 4D**). Additionally, treatment of everted, apical-out spheroids with FAK inhibitor also reduced the infilling of the lumen with cells (**Figure 4C, 4D**). In both cases the FAK inhibitor (pre-treatment or post-treatment) appeared to drive formation of an apical-in actin network; however, podocalyxin staining exhibited an intermittent staining pattern (**Figure 4C, 4D**). Thus we conclude that FAK inhibitors can restore hollow lumens but may only partially restore apical-in polarity.

**Figure 4.**
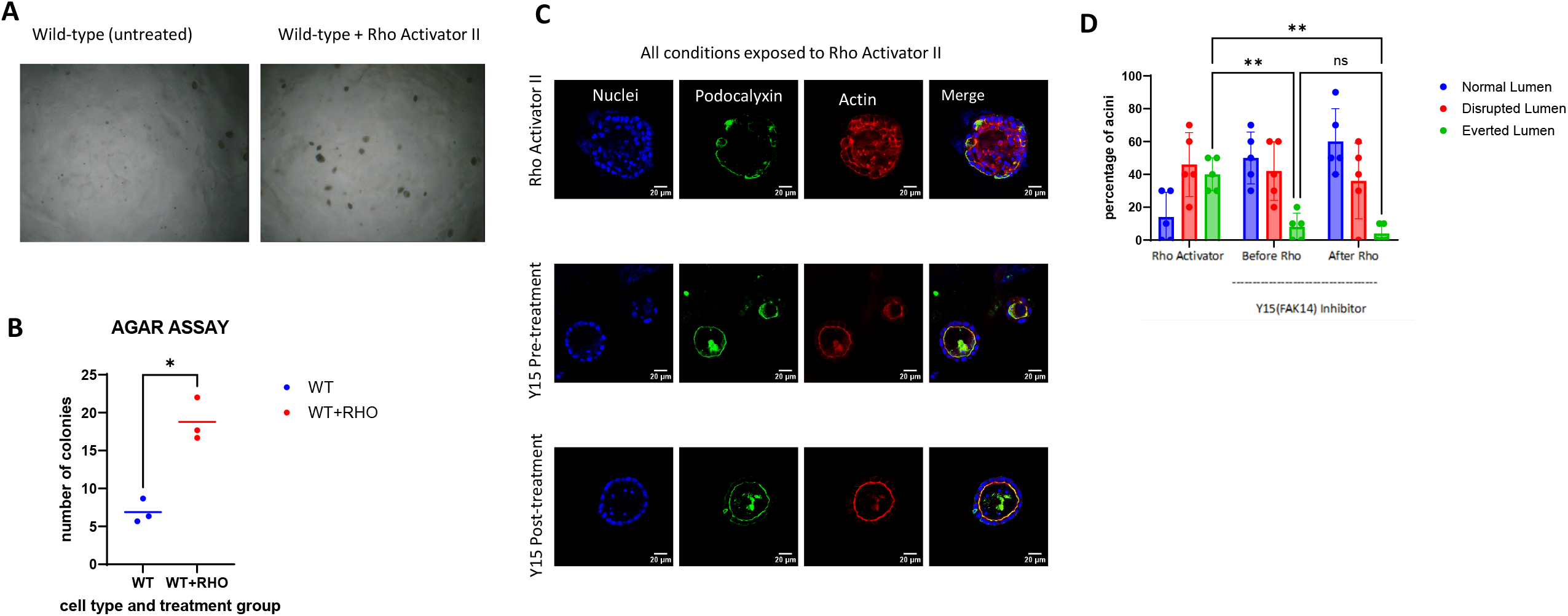
Anchorage-independence is required for apical-out polarity. **A)** MDCK cells were seeded into soft agar. Cells were left untreated or were treated with RhoA activator II. **B)** Quantification of colonies formed in the agar assay. **C)** MDCK cysts were grown in Matrigel for 7 days and then treated with RhoA activator II for 72 hours, pre-treated with FAK inhibitor Y15 24 hours followed by Y15 + Rho Activator II for an additional 48 hours, or treated with Rho Activator II for 24 hours, followed by Y15 + Rho Activator II for an additional 48 hours (post-treatment). **D)** Quantification of normal, disrupted lumen, and eversion for each condition.

### Apical-out polarity is maintained by increased proliferation

Because apical-out spheroids appeared to contain higher number of cells, we hypothesized that apical-out spheroids may have a higher proliferation rate. Using an EdU proliferation assay, we observed that Rho Activator II treatment significantly increased incorporation of EdU into cells, indicating that these spheroids had increased proliferation (**Figure 5A, 5B**). To investigate if RhoA-treatment itself (versus apical-out polarity) is the driver of increased proliferation we also performed the EdU assay in Rho Activator II treated α6 and β4 knockout cells. α6 and β4 knockout cells had a significant increase in cell proliferation following Rho Activator II treatment; however, this increase in proliferation was significantly lower as compared to wild-type cells (**Figure 5A, 5B**). Thus, we conclude that transitions to apical-out polarity further enhance Rho-induced increases in proliferation. Next, we assessed if inhibiting cell proliferation would affect apical-out polarity. Treatment of spheroids with aphidicolin before Rho Activator II treatment or after eversion resulted in spheroids with normal actin polarity and reduced cell infilling (**Figure 5C, 5D**). Similar to FAK inhibition (**Figure 4C**), aphidicolin also resulted in an irregular podocalyxin staining, suggesting that inhibition of polarity only partially restores apical-basal polarity.

**Figure 5.**
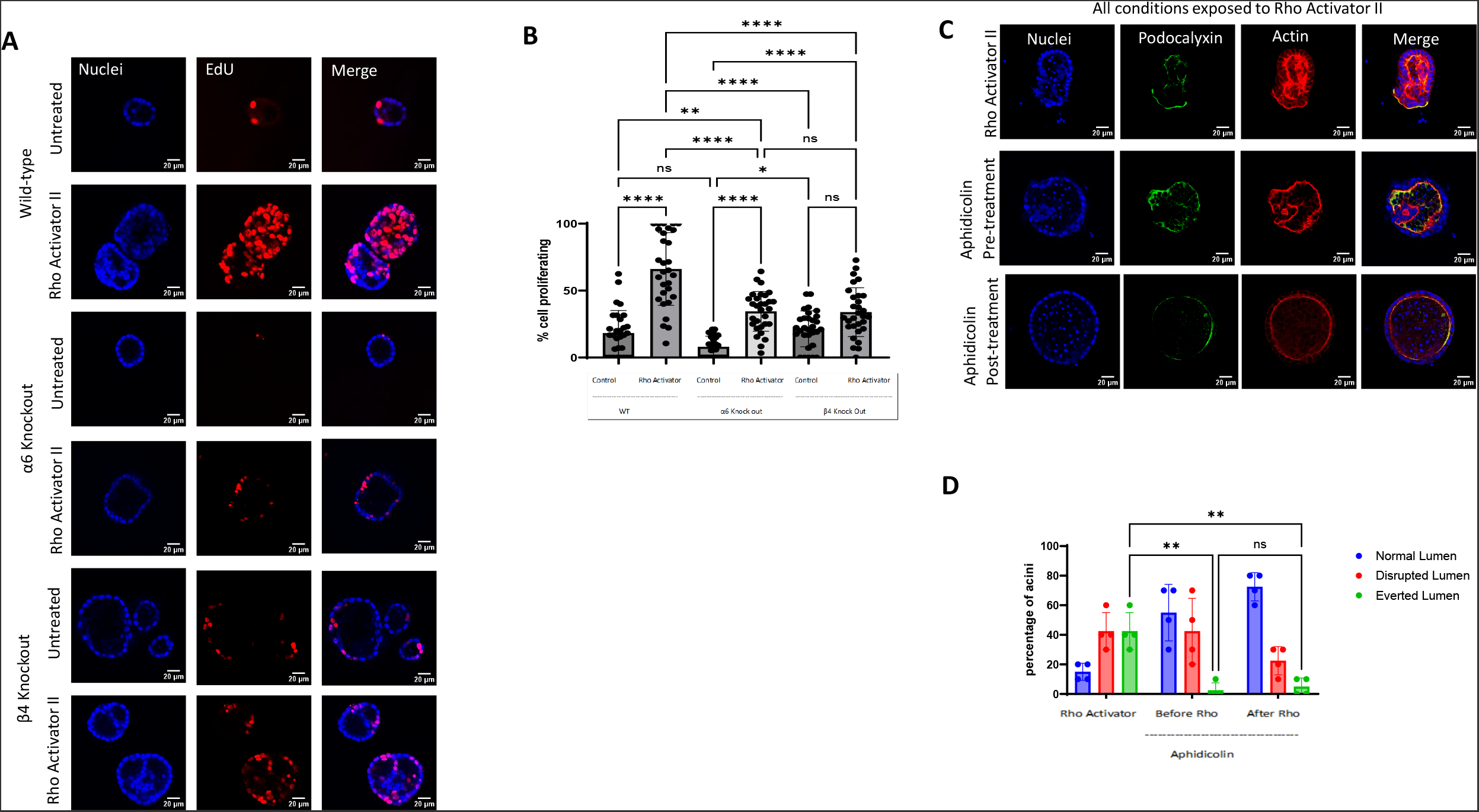
Increased cell proliferation is required to maintain apical-out polarity. **A)** MDCK cells (wild type, α6 integrin knockout, or β4 integrin knockout) were grown in Matrigel for 7 days. Cells were left untreated or treated with RhoA activator II for an additional 48 hours. Cells were incubated with EdU for the final 24 hours of RhoA treatment. Cells were then fixed and EdU incorporation assed using anti-EdU antibodies. **B)** Quantification of the percent of cells with EdU incorporation. **C)** MDCK cysts were grown in Matrigel for 7 days and then treated with RhoA Activator II for 72 hours, pre-treated with DNA synthesis inhibitor aphidicolin for 24 hours followed by aphidicolin + Rho Activator II for an additional 48 hours, or treated with Rho Activator II for 24 hours to induce eversion, followed by aphidicolin + Rho Activator II for an additional 48 hours (post-treatment). **D)** Quantification of normal, disrupted lumen, and eversion for each condition.

### Altered gene expression is observed for apical-out spheroids

We compared the gene expression profiles for wild-type and α6 knockout MDCK cysts, with and without Rho Activator II using bulk RNAseq (**Figure 6A, Supplemental Table 1**). We observed significant changes in gene expression for wild-type cells treated with Rho Activator II (**Figure 6B**). Gene ontology pathways that were significant included proteins associated with extracellular region and growth factor binding (**Figure 6C, Supplemental Table 2**), suggesting that Rho activation and eversion affect how cells interact with the surrounding environment. Given the large number of effects of Rho on gene expression, we wanted to determine which gene expression changes may be most associated with eversion (either causing eversion or downstream of eversion). We compared α6 knockout cells versus wild-type cells each treated with RhoA (**Figure 6D**). Gene ontology pathways which were significantly different between wild-type and α6 knockout cells included pathways related to cellular movement, microtubule movement, extracellular region, chemokine activity, cytokine activity, and GPCR binding (**Figure 6E, Supplemental Table 3**).

**Figure 6.**
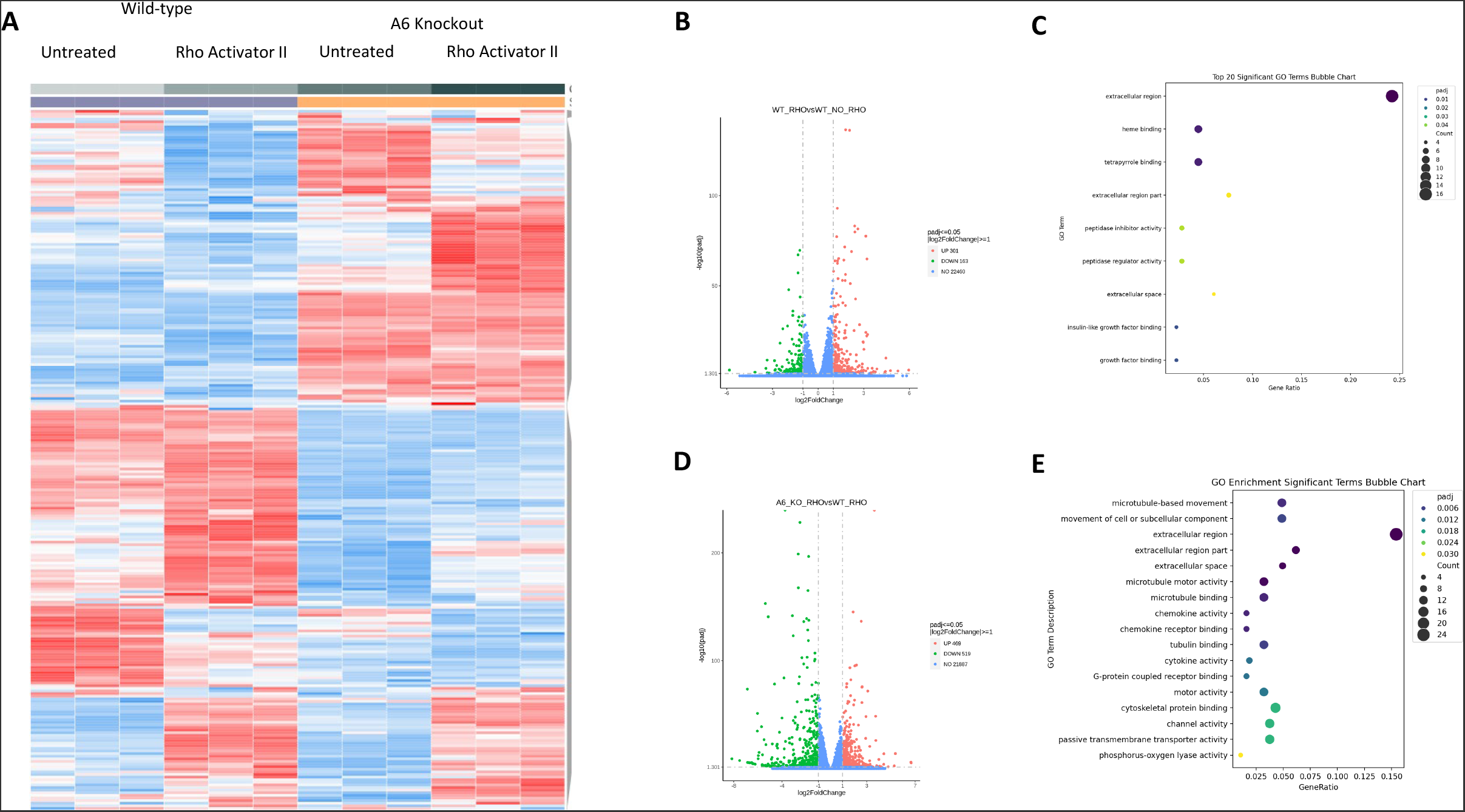
RNA seq analysis of MDCK spheroids. **A)** MDCK spheroids (wildtype or α6 knockout) were left untreated or treated with 48 hours Rho Activator II. Heatmap showing differentially expressed genes (red is upregulated, blue is downregulated). Samples were run in triplicate for each condition. **B)** Volcano plot of wild-type Rho Activator II vs wild-type untreated to show differentially expressed genes from Rho activation. **C)** Gene ontology pathways with significant changes between wild-type Rho Activator and wild-type untreated conditions. **D)** Volcano plot of α6 knockout Rho Activator II vs wild-type Rho Activator II to show differentially expressed genes between everted and non-everted spheroids. **E)** Gene ontology pathways with significant changes between α6 knockout Rho Activator and wild-type Rho Activator II conditions.

## Discussion

Our study provides new mechanistic insights into the mechanisms that drive eversion, by demonstrating the involvement of α6β4 integrins and laminin in promoting eversion (**Figures 1 and 2**). We also demonstrate that apical-out polarity is maintained through increases in cell proliferation and the loss of anchorage-dependence (**Figures 4 and 5**). Interestingly, spheroids with apical-out polarity can revert to apical-in polarity when RhoA activation is stopped (**Figure 3A**), when cell proliferation is inhibited (**Figure 5C**), when spheroids are treated with α6 blocking antibody (**Figure 3D**), or when focal adhesion kinase activity is inhibited (**Figure 4C**), showing that epithelial spheroids can readily transition between apical-in and apical-out polarity. Lastly, we demonstrate that eversion induces significant changes in gene expression (**Figure 6**).

Our study presents the concept that apical polarity is a marker for changes in epithelial cell phenotypes, including increased proliferation and anchorage-independence. It is an open question if apical-out polarity itself drives additional changes to cellular phenotypes. It is tempting to speculate that changes in apical-basal polarity orientation would likely affect the composition of signaling receptors on the outermost surface of the spheroid. If true, we would expect that apical-out spheroids would have different intracellular signaling in response to soluble ligands when compared to spheroids with apical-in polarity.

Apical-out polarity is a pathological feature of specific types of carcinomas, including invasive micropapillary carcinoma (IMPC). Although IMPC was originally identified in breast cancer,^27–30^ IMPC has also been observed in carcinomas of many other organs, including the colon, stomach, rectum, bladder, lung, ovary, and salivary glands, suggesting that apical-out polarity is a reoccurring feature epithelial cancers.^8,31,32^ It is not yet known if eversion is the mechanism that induces apical-out polarity in these cancers—it may be possible that alternate mechanisms, such as altered protein trafficking, establish apical-out polarity *in vivo*. Regardless of the mechanism of establishing apical-out polarity, we postulate that there are parallels between our *in vitro* spheroid model of apical-out polarity and *in vivo* apical-out polarity, where apical-out polarity may be indicative of phenotypic changes in *in vivo* cell behavior. Supporting the concept of an apical-out phenotypic switch, apical-out IMPC is associated with increased metastases to local lymph nodes and lymphovascular invasion, with apical-out polarity most often observed at the invasion front.^8,31,32^ The increased incidence of lymphatic invasion of cancers with reversed apical-basal polarity strongly suggest that cells with apical-out polarity are somehow advantaged for invasion and metastatic spread of cancer. It will be interesting to determine if apical-out polarity in cancer is also correlated to cell phenotypes to those observed in this study of epithelial spheroids (increased cell proliferation, anchorage-independence, altered gene expression).

An important result of our studies is that epithelial cells that have developed apical-out polarity (along with a cell-filled lumen) can readily revert to apical-in polarity (including returning to a hollow cell-free lumen) once the eversion-driving or apical-out-promoting stimuli are eliminated (**Figure 3**). This demonstrates that apical-out polarity is not a permanent state, and that cells can be stimulated to return to normal apical-in polarity. It would be interesting to determine under which conditions would an apical-out epithelial carcinoma return to apical-in polarity, and if this conversion to apical-in polarity could be used as a therapeutic target to reduce or eliminate the invasiveness typically observed in apical-out carcinomas.^8,33^

α6β4 integrins are best known for their role in forming hemidesmosomal adhesions, interacting with intermediate filaments inside the cell and binding extracellularly to laminin. Our work showing that these integrins are important for eversion (**Figure 1**) may be relevant to prior work showing that α6β4-integrins can promote or enhance tumorigenesis^34–36^ and promote anchorage independence.^16,37,38^ Additionally, a prior study showed that oncogenic Ras was shown to upregulate α6 integrin expression, and that α6 integrins promote enhanced metastatic capacity.^16^ It will therefore be interesting to determine if apical-out polarity (through eversion or other mechanisms) is contributing to the enhanced tumorigenesis and metastasis associated with α6β4 upregulation. We also note that α6β4 integrins have been shown to be enriched in K14+ invasive leader cells in breast cancer,^39,40^ although it is not known if or how these integrins contribute to collective metastasis.

Our results demonstrate that the ‘reversion’ from apical-out to apical-in polarity occurs through apoptotic clearance of cells located in the center of the spheroid (**Figure 3C**). We note that this mechanism appears similar if not identical to apoptotic cavitation that has been observed during developmental processes that drive formation of epithelial tubular structures.^41^ Additionally, Mostov and colleagues have shown that apoptotic cavitation is an additional mechanism by which lumen formation occurs in *in vitro* spheroids.^6,42^

Our results also show that the apical-out spheroid stability is promoted in part by increased cell proliferation (**Figure 5**) and anchorage-independence (**Figure 4**). Restoration of anchorage-dependence (through FAK inhibition or treatment with an α6 integrin blocking antibody) or inhibition of proliferation was sufficient to promote clearance of cells from the lumen and a partial return to normal apical-in polarity. We therefore conclude that anchorage-dependent apoptotic clearance is an important process for maintaining a cell-free, hollow lumen. Additionally, expression of oncogenic H-Ras and activation of RhoA (both previously shown stimulate eversion^11^) have each been shown to promote anchorage-independence.^43,44^ Thus is it tempting to speculate that oncogenic acquisition of anchorage-independence is sufficient to promote apical-out polarity in cancer.

In summary our work further elucidates how apical-out polarity is established and is subsequently maintained. In our *in vitro* epithelial model, changes in apical-basal polarity are correlated to phenotypic switching of epithelial cells. It may therefore be notable that the phenotypes we identified associated with apical-out polarity (α6β4 integrins, anchorage-independence, increased cell proliferation) have all been independently associated with cancer progression and invasion.^34–38,45,46^ Future work is needed to determine if carcinomas with apical-out polarity, which is also correlated to cancer invasiveness, ^8,31,32^ have increased anchorage-independence, increased proliferation, or similar gene expression profiles to everted spheroids.

## Acknowledgements

We acknowledge helpful discussions with Richard Dickinson, Alex Dunn, Andrew Ewald, Priscilla Hwang, Jessica Williamson, and Tanmay Lele. We also wish to thank Gillian DeWane and Kris DeMali for advice on optimization of the soft agar assay and Scott Crawley for advice on immunostaining the apical membrane in Caco-2 cells. This work was funded by NIH grant R35 GM119617.

## Figure Legends

**Supplemental Figure 1. Screening of additional MDCK integrin knockout lines. A)** β1 knockout and β1+β4 double knockout MDCK cells were seeded in Matrigel and assessed for lumen formation and apical-basal polarity. These cells formed spheroids with apical-out polarity. **B)** α3 knockout and αV knockout cells were seeded in Matrigel and assessed for lumen formation and apical-basal polarity. These spheroids exhibited a multi-luminated phenotype. Treatment with 48 hours of Rho A activator II resulted in apical-out polarity.

**Supplemental Figure 2. α6 blocking antibody treatment has minimal effects in unstimulated MDCK spheroids**. MDCK spheroids were treated with α6 blocking antibody. Cysts were fixed and immunostained for phospho-Ezrin and actin (phalloidin). In some instances no disruption in apical-basal polarity was observed (top), whereas for other spheroids podocalyxin was observed on both the inner and outer membranes (bottom).

**Supplemental Figure 3. Eversion of Caco-2 cells requires α6 integrins. A)** Caco-2 cells were seeded in a mixture of collagen and Matrigel. Mature cysts were left untreated, or treated with α6 blocking antibody, RhoA activator II, or α6 blocking antibody with RhoA activator II. Cysts were fixed and immunostained for phospho-Ezrin and actin (phalloidin). **B)** Quantification of normal, disrupted lumen, and eversion for blocking antibody experiments.

**Supplemental Table 1. Processed RNAseq data**

**Supplemental Table 2. Gene ontology pathway analysis for wild-type untreated vs wild-type Rho Activator II treated spheroids.**

**Supplemental Table 3. Gene ontology pathway analysis for α6 knockout Rho Activator II treated spheroids vs wild-type Rho Activator II treated spheroids.**

