## Supplemental Figures 1-3 for "Epithelial eversion, a collective rearrangement from apical-in to apical-out polarity, is initiated by α6β4 integrins and sustained by increased cell proliferation and anchorage-independence"

**Supplemental Figure 1. Screening of additional MDCK integrin knockout lines. A)** β1 knockout and β1+β4 double knockout MDCK cells were seeded in Matrigel and assessed for lumen formation and apical-basal polarity. These cells formed spheroids with apical-out polarity. **B)** α3 knockout and αV knockout cells were seeded in Matrigel and assessed for lumen formation and apical-basal polarity. These spheroids exhibited a multi-luminated phenotype. Treatment with 48 hours of Rho A activator II resulted in apical-out polarity.

**Supplemental Figure 2. α6 blocking antibody treatment has minimal effects in unstimulated MDCK spheroids.** MDCK spheroids were treated with α6 blocking antibody. Cysts were fixed and immunostained for phospho-Ezrin and actin (phalloidin). In some instances no disruption in apical-basal polarity was observed (top), whereas for other spheroids podocalyxin was observed on both the inner and outer membranes (bottom).

**Supplemental Figure 3. Eversion of Caco-2 cells requires α6 integrins. A)** Caco-2 cells were seeded in a mixture of collagen and Matrigel. Mature cysts were left untreated, or treated with α6 blocking antibody, RhoA activator II, or α6 blocking antibody with RhoA activator II. Cysts were fixed and immunostained for phospho-Ezrin and actin (phalloidin). **B)** Quantification of normal, disrupted lumen, and eversion for blocking antibody experiments.

**Supplemental Table 1. Processed RNAseq data**

**Supplemental Table 2. Gene ontology pathway analysis for wild-type untreated vs wild-type Rho Activator II treated spheroids.**

**Supplemental Table 3. Gene ontology pathway analysis for α6 knockout Rho Activator II treated spheroids vs wild-type Rho Activator II treated spheroids.**

### Supplemental Figure 1

A

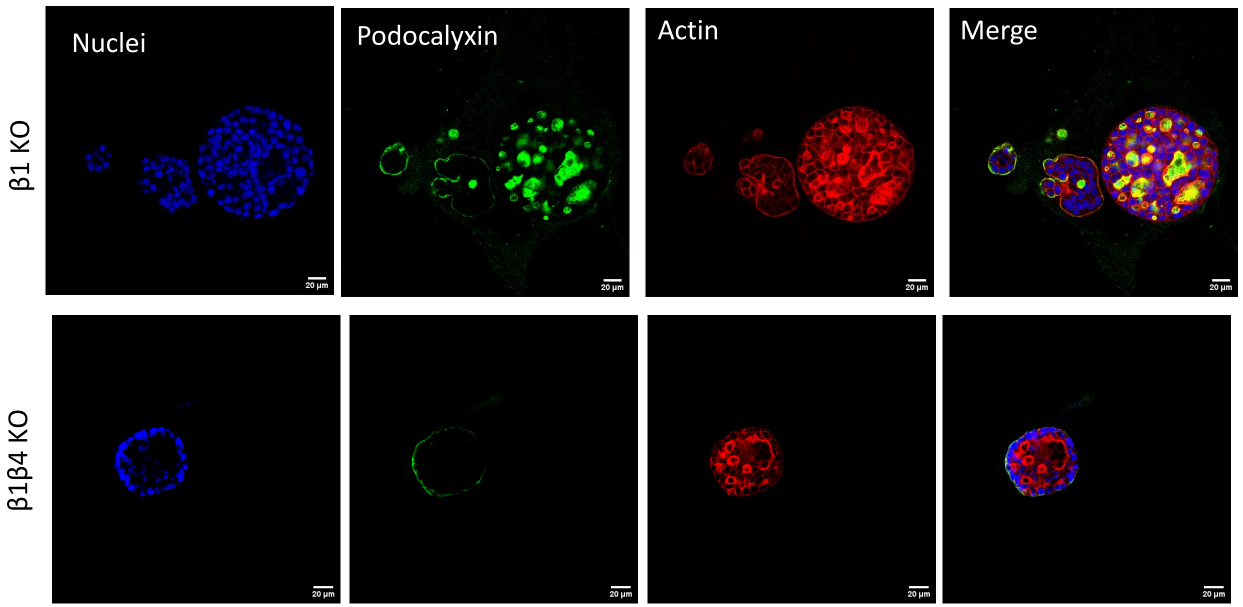


B

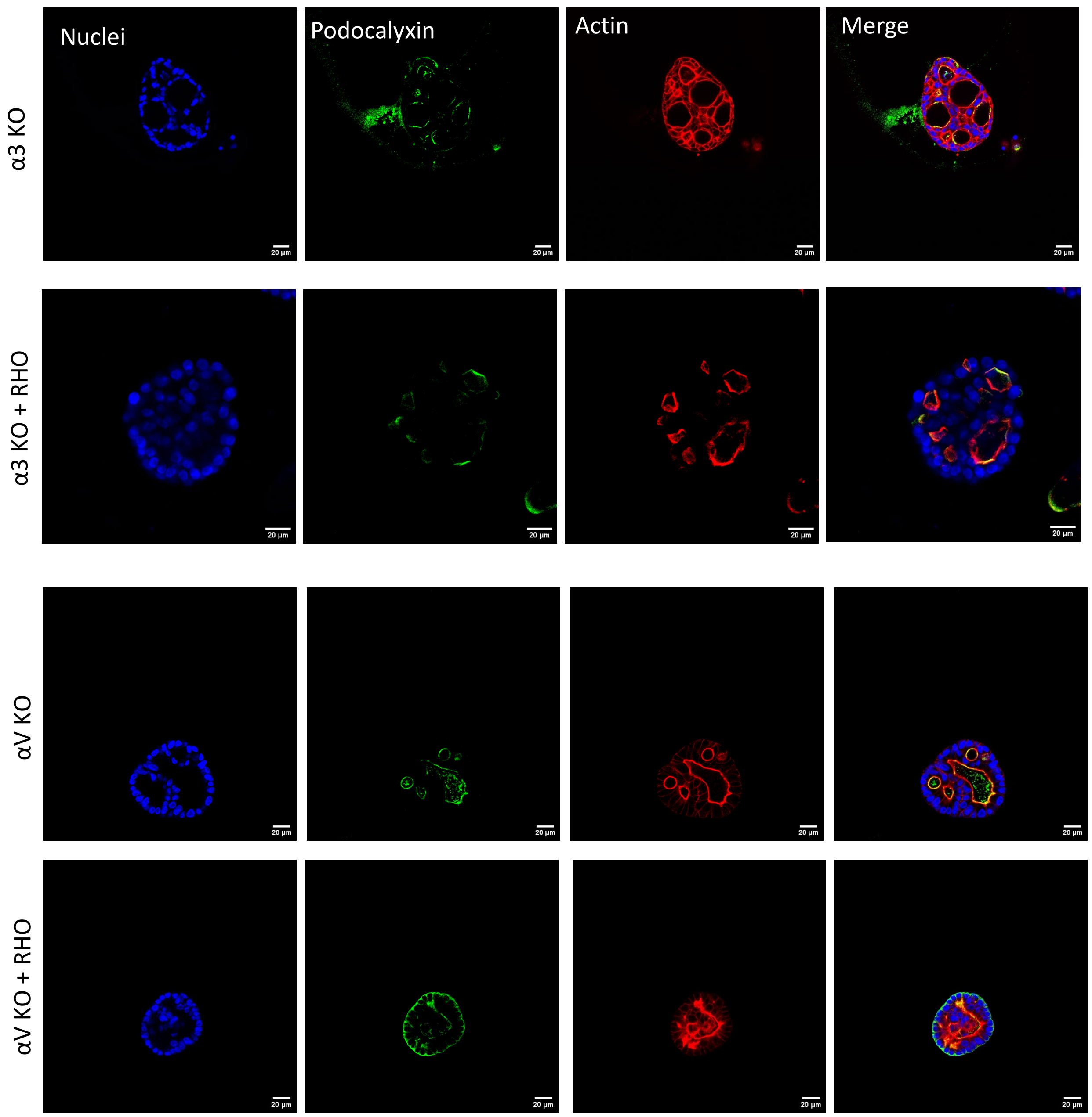

### Supplemental Figure 2

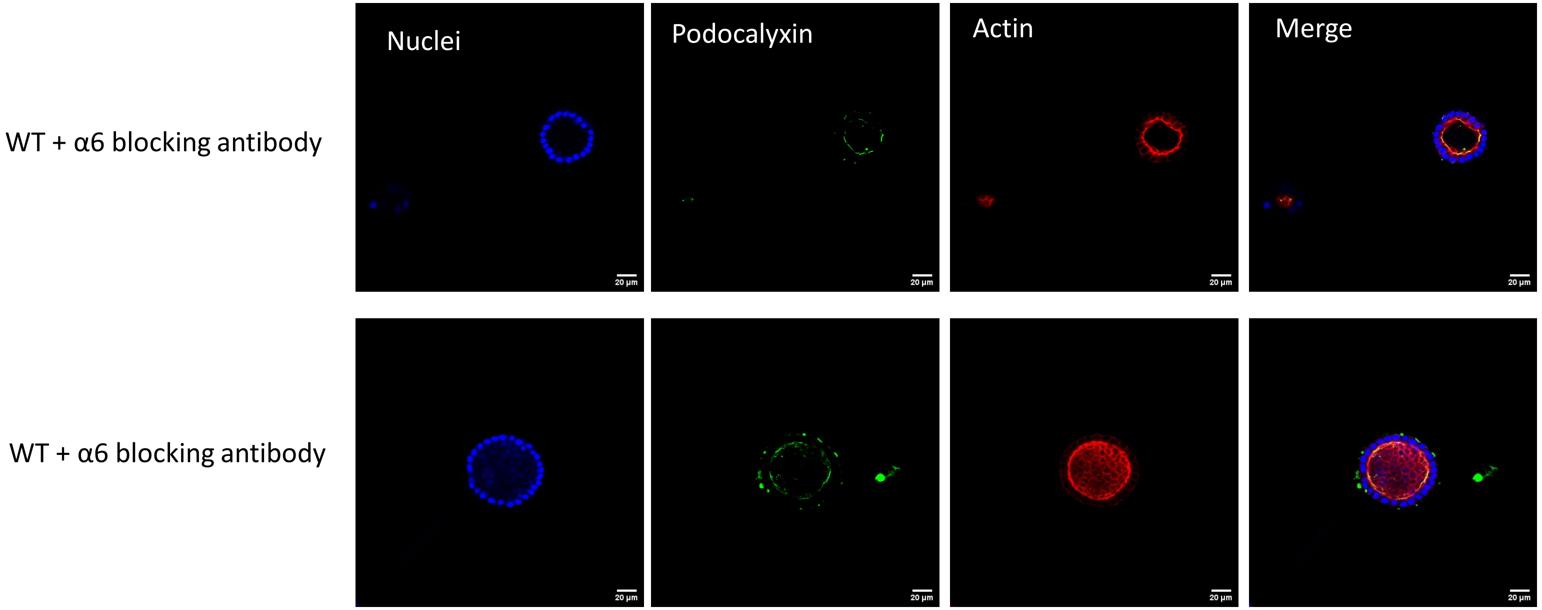

### Supplemental Figure 3

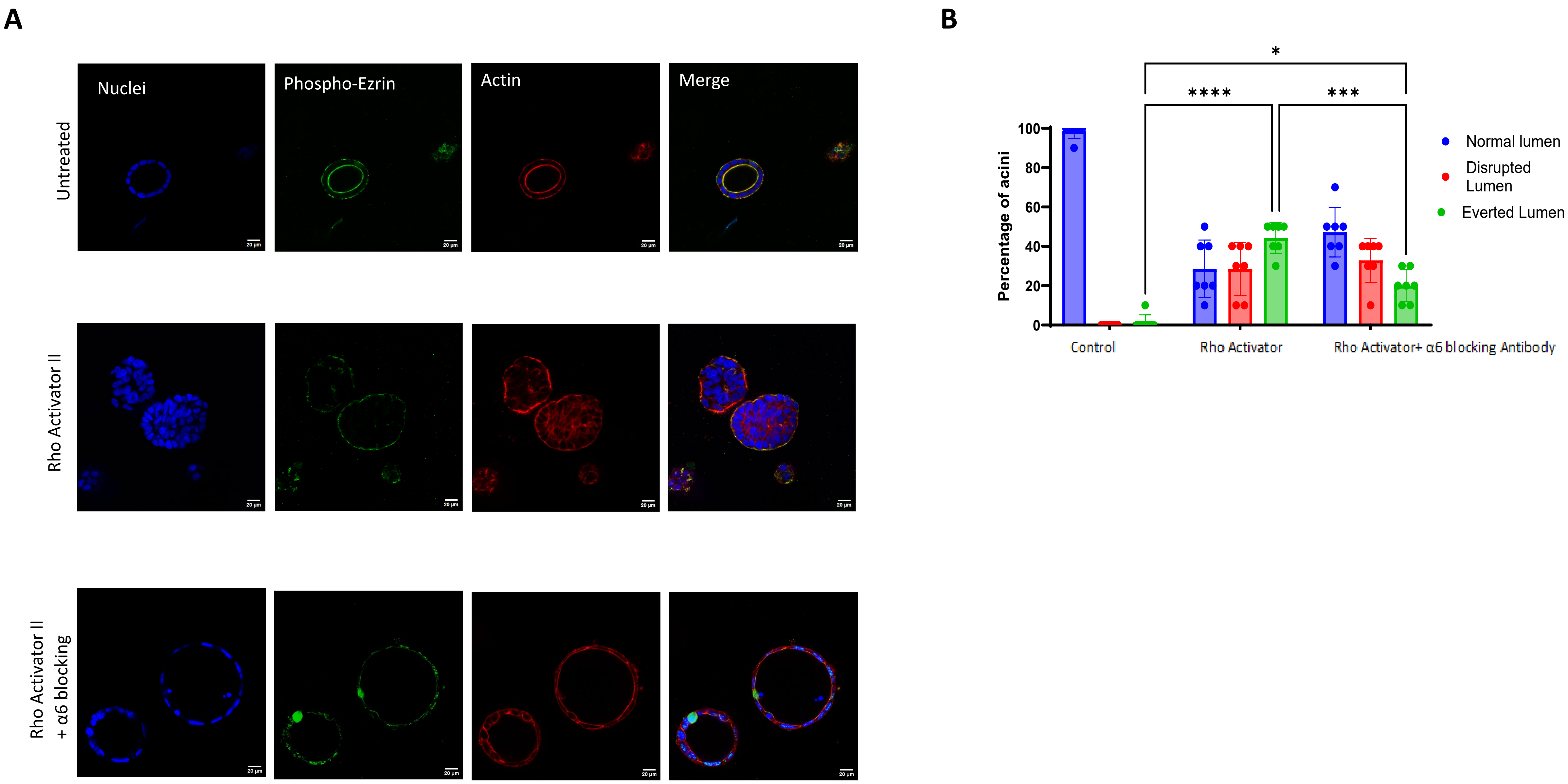
